# Angiogenic mechanisms governing the segregation of blood-brain barrier and fenestrated capillaries derived from a multipotent cerebrovascular niche

**DOI:** 10.1101/2024.12.10.627641

**Authors:** Nathanael J. Lee, Sweta Parab, Amanda E. Lam, Jun Xiong Leong, Ryota L. Matsuoka

## Abstract

Cerebrovascular endothelial cell (EC) subtypes characterized by blood-brain barrier (BBB) properties or fenestrated pores are essential components of brain-blood interfaces, supporting brain function and homeostasis. To date, the origins and developmental mechanisms underlying this heterogeneous EC network remain largely unclear. Using single-cell-resolution lineage tracing in zebrafish, we discover a multipotent vascular niche at embryonic capillary borders that generates ECs with BBB or fenestrated molecular identity. RNAscope analysis demonstrates restricted expression of *flt4* in sprouting ECs contributing to fenestrated choroid plexus (CP) vasculature, identifying an early molecular distinction from adjacent BBB vessels. Mechanistically, *flt4* null and cytoplasmic-domain-deletion mutants exhibit CP vascularization defects when combined with *vegfr2* zebrafish paralog deletion. Pharmacological results support this co-requirement of Flt4 and Vegfr2 signaling and suggest the PI3K and ERK pathways as downstream effectors. These findings reveal a specialized developmental origin for BBB and fenestrated EC subtypes, and establish Flt4 as a crucial guidance receptor mediating their angiogenic segregation.

## INTRODUCTION

Striking differences in cerebrovascular permeability were first observed in the early 1900s, when studies showed that intravenous injections of trypan blue strongly stained the CP, but not the brain parenchyma (*1, 2*). Subsequent research has characterized cerebrovascular barrier and non-barrier properties at anatomical and functional levels. Current evidence suggests that cerebrovascular ECs play a central role in region-specific vascular permeability through their phenotypic differences (*3, 4*). These differences are most evident at the capillary level, where ECs establish the semi-permeable BBB in most regions but form permeable fenestrations in the CPs and circumventricular organs (CVOs) (*5, 6*). Both EC subtypes are essential for maintaining the specialized neural environment, yet the developmental processes governing this capillary segregation remain poorly understood.

Like other peripheral organs, brain vascularization relies on angiogenesis. Blood vessel invasion into the brain parenchyma begins with sprouting angiogenesis from the perineural vascular plexus (*7, 8*). This process requires tip cell specialization dependent on Wnt7/β-catenin signaling and its downstream effector, Mmp25 (*9, 10*), where Gpr124 and Reck serve as receptor co-factors crucial for β-catenin activation (*10–12*). Subsequently, endothelial β-catenin drives gene transcription critical for establishing BBB properties (*13–15*). These angiogenic and BBB induction mechanisms are evolutionarily conserved between mammals and zebrafish (*6, 16*).

Conversely, much less is known about the developmental mechanisms for fenestrated capillary formation. Our recent studies suggest that Vegfa and Vegfc/d ligands are co-required for angiogenesis leading to CP and CVO vascularization in zebrafish, with this requirement fulfilled locally through interactions between growing ECs and specialized cell types in these organs (*17–19*). Interestingly, despite the severe loss of CP vasculature in the absence of these Vegfs, the formation of neighboring BBB vessels remains largely unaffected (*18*). This striking specificity raises the question of what molecular mechanisms enable EC’s differential responsiveness to locally presented angiogenic cues.

Here we investigated underlying mechanisms of this specificity and unexpectedly uncovered a cerebrovascular niche that generates both CP and BBB EC subtypes. Our mechanistic studies further identified key receptors and downstream effectors driving their angiogenic segregation.

## RESULTS

### Single-cell-resolution lineage tracing identifies a cerebrovascular niche that generates ECs contributing to fenestrated CP vasculature and adjacent BBB vessels

Across vertebrates, the CPs maintain brain homeostasis by producing cerebrospinal fluid and clearing metabolic wastes (*20*). In mammals, CPs develop on the surfaces of the lateral, 3rd, and 4th brain ventricles (*21*), but their deep anatomical locations and *in-utero* development present significant challenges for live imaging. To address this limitation, we leveraged the zebrafish model. The zebrafish CPs form in two anatomical locations: the diencephalic and myelencephalic CP (dCP and mCP, respectively) (*22*). Both CPs are located near the dorsal brain surfaces, enabling non-invasive visualization of CP vascular morphogenesis at single-cell resolution. The mCP is larger and positioned near the 4th ventricle and cerebellum (Fig. 1A).

**Fig. 1.**
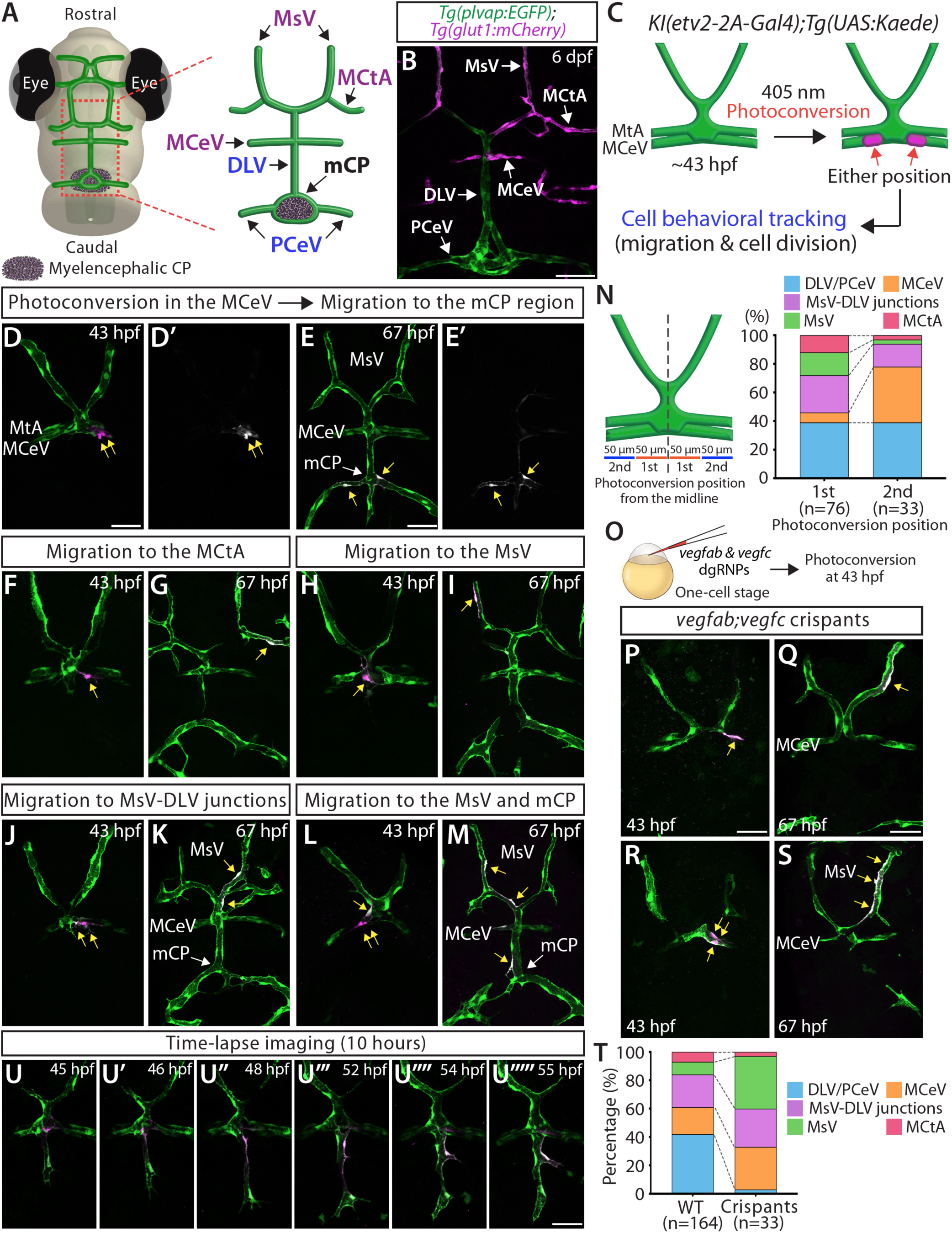
Identification of a specialized cerebrovascular niche generating ECs that migrate to diverse blood vessels. (**A**) Diagram of the larval zebrafish head, indicating the location of the mCP and associated blood vessels: DLV, PCeV, MsV, MCeV, and MCtA display distinct molecular signatures. (**B**) Dorsal view of 6 dpf *Tg(plvap:EGFP);Tg(glut1:mCherry)* larval head, showing strong *plvap* expression in the DLV and PCeV, and restricted *glut1* expression in the MsV, MCeV, and MCtA. (**C**) Experimental setup for photoconversion experiments (**D**-**N**). (**D**-**E’**) Photoconversion in 43 hpf *KI(etv2-2A-Gal4);Tg(UAS:Kaede)* embryo (**D**, **D’**), with subsequent tracking at 67 hpf (**E**, **E’**). Photoconverted MCeV cells migrated to the mCP region (yellow arrows). (**F**-**M**) Examples of photoconverted ECs in the MCeV, migrating to various vessels. Photoconversion performed at 43 hpf (**F**, **H**, **J**, **L**) with subsequent tracking at 67 hpf (**G**, **I**, **K**, **M**). Photoconverted ECs migrated to the MCtA (**G**), MsV (**I**, **M**), MsV-DLV junctions (**K**, **M**), and DLV (**M**). (**N**) Summary of photoconversion experiments in the MCeV at 43 hpf, showing diverse migration destinations of ECs. Initial photoconversion positions were categorized by their distance from the midline, as illustrated. Graphs include data from the 1st position (n=76) and the 2nd position (n=33). (**O**) Experimental setup for microinjection experiments (**P**-**T**). (**P**-**S**) Photoconversion in *vegfab;vegfc* crispants at 43 hpf (**P** and **R**) and 67 hpf (**Q** and **S**). These crispants lacked the DLV, and photoconverted ECs migrated to the MsV (yellow arrows). (**T**) Graphs comparing all combined photoconversion results between WT (n=164) and *vegfab;vegfc* crispants (n=33). These graphs include similar proportions of ECs photoconverted from the 1st position (62% for WT and 61% for *vegfab;vegfc* crispants). (**U**-**U””’**) Time-lapse image from 45 hpf to 55 hpf, showing photoconverted cell migration to the DLV. Scale bars: 50 µm in **B**; 50 µm in **D** for **F**, **H**, **J**, **L**; 50 µm in **E** for **G**, **I**, **K**, **M**; **O**; 50 µm in **P** for **R**; 50 µm in **Q** for **S**; 50 µm in **U** for **U**-**U””’**.

Current evidence suggests that mCP vasculature sprouts from the middle cerebral vein (MCeV) (*23*) and that the MCeV acquires BBB properties, while the mCP vasculature exhibits fenestrated features (Fig. 1B) (*18, 24*). An important developmental question remains unanswered: how do these vessels segregate to establish the heterogeneous vascular networks? To address this question, we investigated early events that drive mCP vessel sprouting from the MCeV in zebrafish embryos.

Consistent with previous findings, we observed that the MCeV forms near the metencephalic artery (MtA) at the midbrain-hindbrain boundary, residing caudally between 1.5 and 2 days post fertilization (dpf) (Fig. S1S) (*25*). During this period, the dorsal longitudinal vein (DLV) initiates sprouting to form mCP vasculature by fusing with bilateral posterior cerebral vein (PCeV) through 75 hours post fertilization (hpf) (Fig. S1T-U). From this stage onward, the MCeV and MtA become intermingled, appearing as a single vessel that is designated as the MCeV (Fig. S1U-V).

While a previous study indicated DLV sprouting from the MCeV in a pan-EC transgenic line (*23*), its limited resolution made it difficult to pinpoint specific ECs contributing to the mCP vasculature. To address this issue, we applied EC-specific photoconversion to trace EC lineages. Specifically, we employed the knock-in line, herein *KI(etv2-2A-Gal4)*, which drives Gal4 expression under the endogenous promoter of the early angioblast marker *etv2* (*26*). Crossing this line with *Tg(UAS:Kaede)* adults produced progeny expressing the photoconvertible Kaede protein specifically in ECs.

We photoconverted Kaede from green to red fluorescence in specific ECs by applying 405 nm laser illumination for 1-2 seconds (Fig. 1C). Using this approach, we began by targeting a few ECs on the MCeV at approximately 43 hpf before DLV sprouting initiated, and subsequently tracked these cells 24 hours later at 67 hpf. By targeting ECs within various areas of the MCeV, we identified a specific region as the source of ECs contributing to the mCP vasculature (Fig. 1D-E’). This region is located near the dorsal midline junction (DMJ), where DLV sprouting begins (*23*).

While characterizing this area, we surprisingly found that ECs originating from this place migrated to diverse vessels, including those acquiring BBB molecular signatures. Our careful tracking of photoconverted ECs from the MCeV revealed five distinct destinations: DLV/PCeV (mCP vessels) (Fig. 1D-E’), mesencephalic central artery (MCtA) (Fig. 1F-G), mesencephalic vein (MsV) (Fig. 1H-I), MsV-DLV junctions (Fig. 1J-K), and MCeV (Fig. S1J-L). Strikingly, adjacent ECs within this niche often exhibited entirely distinct migration patterns (Fig. 1L-M, S1G-I). To quantify these observations, we categorized photoconverted ECs based on their initial positions relative to the midline. We found that ECs located within 50 µm from the midline (1st position) most preferentially migrated to mCP vessels (39%; Fig. 1D-E’, S1A-F), with the second most frequent destination being the MsV-DLV junctions (26%; Fig. 1N). Nearly 65% of cells from this position migrated to blood vessels with fenestrated features, though MsV-DLV junctions often displayed BBB marker expression on one side (Fig. 1B). The remaining 35% migrated to the MsV (16%), MCtA (12%), and MCeV (7%) (Fig. 1N).

Further analysis of ECs located 50 to 100 µm from the midline (2nd position) revealed a markedly higher proportion of ECs remaining on the MCeV (39%, Fig. 1N). Accordingly, the overall proportions of ECs migrating to other vessels decreased, while migration to mCP vessels remained relatively high at 39%. As a general observation, cells originating on the left side tended to remain on the left, while those starting on the right side largely stayed on the right (Fig. S1A-F, S1J-O).

### The MCeV is a specialized brain vessel bearing multipotent angioblasts during development

To investigate whether the MCeV serves as the specific source of ECs contributing to mCP and nearby BBB vessels, we targeted the adjacent MtA at similar locations relative to the midline (Fig. S1M-O). Intriguingly, ECs on the MtA almost exclusively remained on this vessel (9 out of 10 cells, with the remaining cell migrating to the MCtA), indicating the MCeV as a specialized vessel bearing multipotent angioblasts that generate ECs of fenestrated and BBB identities.

Next, we monitored sprouting angiogenesis at this capillary border using time-lapse imaging with and without photoconversion. During the recorded period from 32 hpf to 47 hpf, we found dynamic EC scrambling near the DMJ, indicative of active exploratory behaviors, both before and during DLV sprouting (Movie S1). Live imaging after photoconversion recorded from 43 to 53 hpf captured progressive migration of targeted EC to diverse vessels, including the DLV (Fig. 1U-U””’, Movie S2). Notably, none of the photoconverted ECs (n=19) underwent cell division during the recorded time period, suggesting that EC migration is a central mechanism driving angiogenesis from this specialized niche.

To explore how ECs migrate to diverse vessels from this niche, we examined an established BBB marker, *Tg(glut1:*mCherry) expression, around the DMJ (Fig. S1S-V). This analysis revealed that the DMJ region is populated by molecularly heterogeneous ECs before DLV sprouting: some ECs exhibited *Tg(glut1:*mCherry) expression, while others did not (Fig. S1W-W”). This observation implies that ECs in this region are pre-determined and primed to migrate in response to specific angiogenic cues.

### The combined loss of Vegfab and Vegfc causes DLV loss, resulting in a biased EC migration from the MCeV toward BBB vessels

Our recent studies showed that the combined loss of Vegfab and either Vegfc or Vegfd (or both) substantially increased DLV-absent phenotypes (*18*). To investigate how the deletion of these factors affects EC behaviors, we employed a CRISPR-based approach that enables efficient biallelic gene inactivation in F0 zebrafish (*27*). Specifically, we performed simultaneous injections of three distinct dual-guide CRISPR RNA/Cas9 ribonucleoproteins (dgRNPs) per gene, which achieved highly consistent generations of F0 knockouts (crispants) (*27*). By co-injecting a cocktail of three dgRNPs targeting *vegfab* and three targeting *vegfc* at the one-cell stage (Fig. 1O), we successfully recapitulated the DLV loss phenotypes observed in stable *vegfab;vegfc* double mutants (*18*). Next, we injected this dgRNP cocktail into one-cell stage embryos carrying both the *KI(etv2-2A-Gal4)* and *Tg(UAS:Kaede)* transgenes to perform photoconversion. We targeted comparable MCeV areas in crispants and tracked photoconverted cells over the same time course as wild-type (WT) embryos. Notably, crispants showed a sharp reduction in cells migrating to mCP vessels and a dramatic increase in cells migrating to the MsV (Fig. 1P-S, 1T, S1P-R). These results suggest that Vegfab and Vegfc co-deletion caused dramatic shifts in EC migration destinations from mCP vasculature to BBB-associated vessels.

### PI3K and MEK signaling pathways mediate Vegfab/Vegfc-dependent mCP vascularization

VEGF ligands regulate angiogenesis by activating downstream kinases, including PI3K, ERK, and PLCK (*28*). To explore the EC intrinsic mechanisms driving migration from this niche, we first focused on identifying intracellular signaling pathways mediating Vegfs’ angiogenic effects. Since the major kinases activated by Vegfs regulate various developmental processes, we employed a pharmacological approach to block individual pathways during a defined time window.

Specifically, we treated embryos between 34 and 72 hpf with the following chemicals: SL327, a MEK1/2 inhibitor (*29*); LY294002, a selective PI3K inhibitor (*30*); and U73122, a PLCK inhibitor (*31*). We first determined sublethal doses of each inhibitor that did not cause gross morphological changes (Fig. 2A-E). Treatment with 2 μM SL327 or 17.5 μM LY294002 significantly increased the DLV-absent phenotype (nearly 40% and 50%, respectively) (Fig. 2F-H, 2J), shortened DLV lengths (Fig. 2K), and impaired PCeV formation (Fig. 2L). In contrast, 1 μM U73122 treatment did not cause any of these vessel formation defects (Fig. 2I-L). These results indicate that the PI3K and MEK pathways, but not PLCK, are activated by Vegfs during DLV and PCeV formation.

**Fig. 2.**
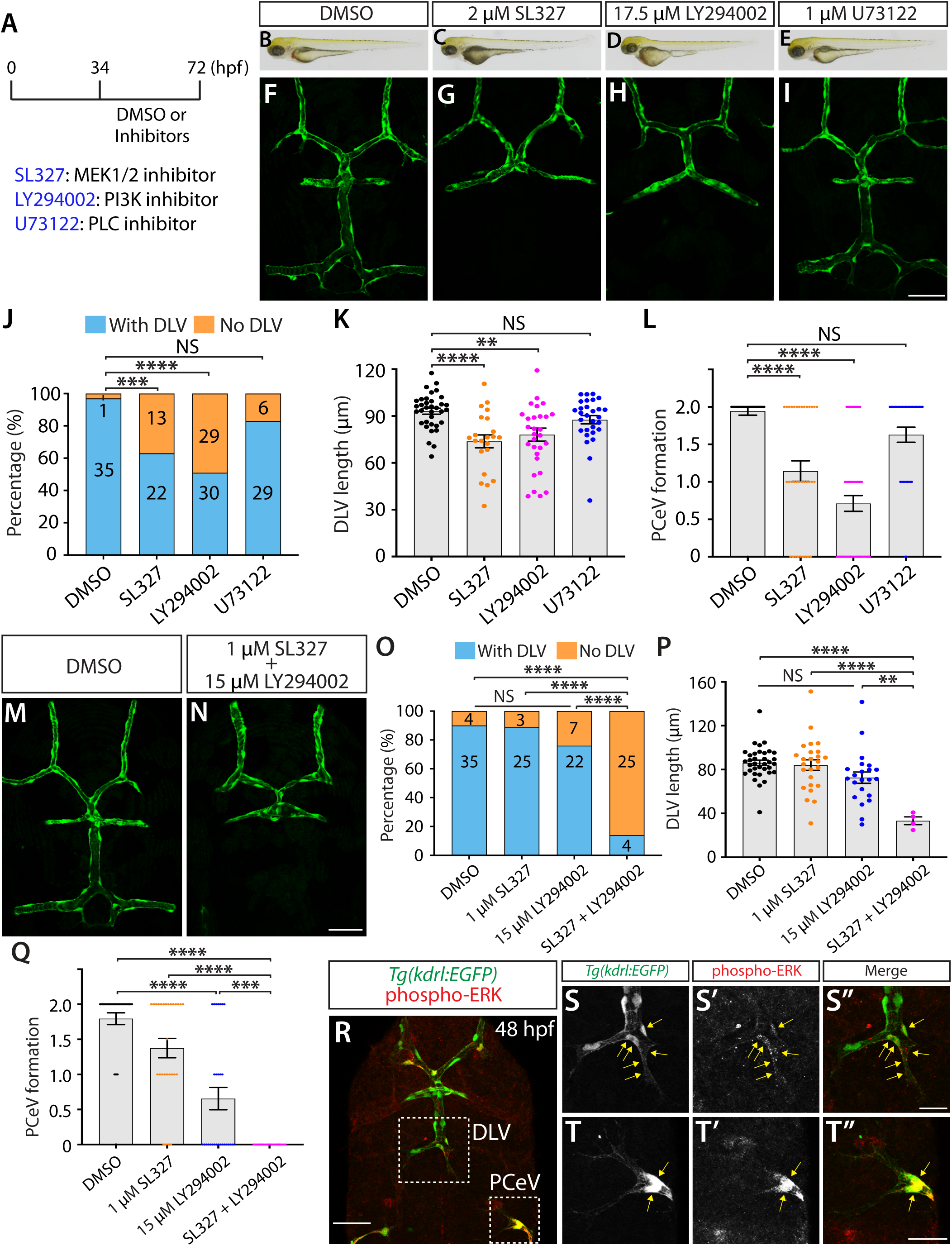
Chemical inhibition of MEK and PI3K signaling disrupted mCP vessel formation. (**A**) Experimental timeline for chemical treatments (**B**-**L**). (**B**-**E**) Brightfield images of 72 hpf larvae after treatment with DMSO (**B**), 2 µM SL327 (**C**), 17.5 µM LY294002 (**D**), and 1 µM U73122 (**E**). (**F**-**I**) Dorsal views of the 72 hpf *Tg(kdrl:EGFP)* larval head after treatment with DMSO (**F**), 2 µM SL327 (**G**), 17.5 µM LY294002 (**H**), and 1 µM U73122 (**I**). (**J**) Percentage of 72 hpf *Tg(kdrl:EGFP)* fish with and without the DLV after treatment (number of animals examined per treatment is listed). (**K**) DLV length quantification in 72 hpf *Tg(kdrl:EGFP)* larvae that formed the DLV (n=35 for DMSO, n=22 for SL327, n=30 for LY294002, and n=29 for U73122). (**L**) Quantification of PCeV formation in 72 hpf *Tg(kdrl:*EGFP) larvae after treatment (n=36 for DMSO, n=35 for SL327, n=59 for LY294002, and n=34 for U73122). (**M** and **N**) 72 hpf *Tg(kdrl:EGFP)* larval head after treatment with DMSO (**M**) and a combined solution of 1 µM SL327 and 15 µM LY294002 (**N**). (**O**) Percentage of 72 hpf *Tg(kdrl:EGFP)* larvae with and without the DLV after treatment (number of animals examined per treatment is listed). (**P**) DLV length quantification in 72 hpf *Tg(kdrl:EGFP)* larvae that formed the DLV (n=35 for DMSO, n=25 for 1 µM SL327, n=22 for 15 µM LY294002, and n=4 for combined SL327/LY294002). (**Q**) Quantification of PCeV formation in 72 hpf *Tg(kdrl:*EGFP) larvae after treatment (n=39 for DMSO, n=28 for 1 µM SL327, n=29 for 15 µM LY294002, and n=29 for combined SL327/LY294002). (**R**) 48 hpf *Tg(kdrl:EGFP)* embryos immunostained for EGFP and phosphorylated ERK. Magnified images of the DLV from (**R**) are shown in (**S**-**S”**) and those of the PCeV in (**T**-**T”**). Scale bars: 50 µm in **I** for **F**-**I**; 50 µm in **N** for **M**; 50 µm in **R**; 20 µm in **S”** for **S**-**S”**; 20 µm in **T”** for **T**-**T”**.

To investigate potential additive or synergistic effects of co-treating with LY294002 and SL327, we used lower concentrations of each inhibitor (1 μM SL327 and 15 μM LY294002) to better detect synergy. At these dosages, individual treatments caused only mild effects on DLV formation, reflected by DLV loss phenotypes (Fig. 2O) with no significant impact on DLV length (Fig. 2P). However, this co-treatment substantially increased defects in DLV and PCeV formation (86%) (Fig. 2M-Q). Similar results were obtained with half the dosage of these combined inhibitors (Fig. S2), suggesting that PI3K and MEK pathways are key effectors in mCP vascularization.

### Phosphorylated ERK is detected in tip cells of the developing DLV and PCeV

If PI3K and MEK pathways are required for mCP vascularization in ECs, their activation should be detectable in growing vessels during DLV and PCeV formation. To test this idea, we performed wholemount immunostaining with an antibody specific to phosphorylated ERK (pERK), a critical activation step in the MAPK signaling cascade. At 50 hpf, when both the DLV and PCeV were extending, we detected strong pERK signals in leading cells of both vessels (Fig. 2R-T”). This result suggests that MEK/ERK signaling is active during mCP vascularization, supporting our pharmacological findings.

### RNAscope analysis identifies *flt4* as an early molecular marker distinguishing mCP vasculature from adjacent BBB vessels

To next identify which Vegf receptors (Vegfrs) contribute to mCP vascularization, we examined *vegfr* gene expression using RNAscope *in situ* hybridization (ISH) (*32*). This method generates specific punctate signals representing individual mRNA transcripts (*32*), enabling spatial mapping of gene expression at single-cell resolution with quantifiable data.

In zebrafish, the Vegfr family includes four members: Flt1, Kdr, Kdrl, and Flt4 (Fig. 3A) (*33*). Biochemical studies showed that Vegfc binds to Flt4 and Kdr, whereas Vegfd binds exclusively to Kdr (*33*). Vegfa paralogs (Vegfaa and Vegfab) interact with Kdr and Kdrl (Vegfr2 paralogs) (*34*), and Flt1 acts as a decoy receptor modulating Vegfa signaling (*35*).

**Fig. 3.**
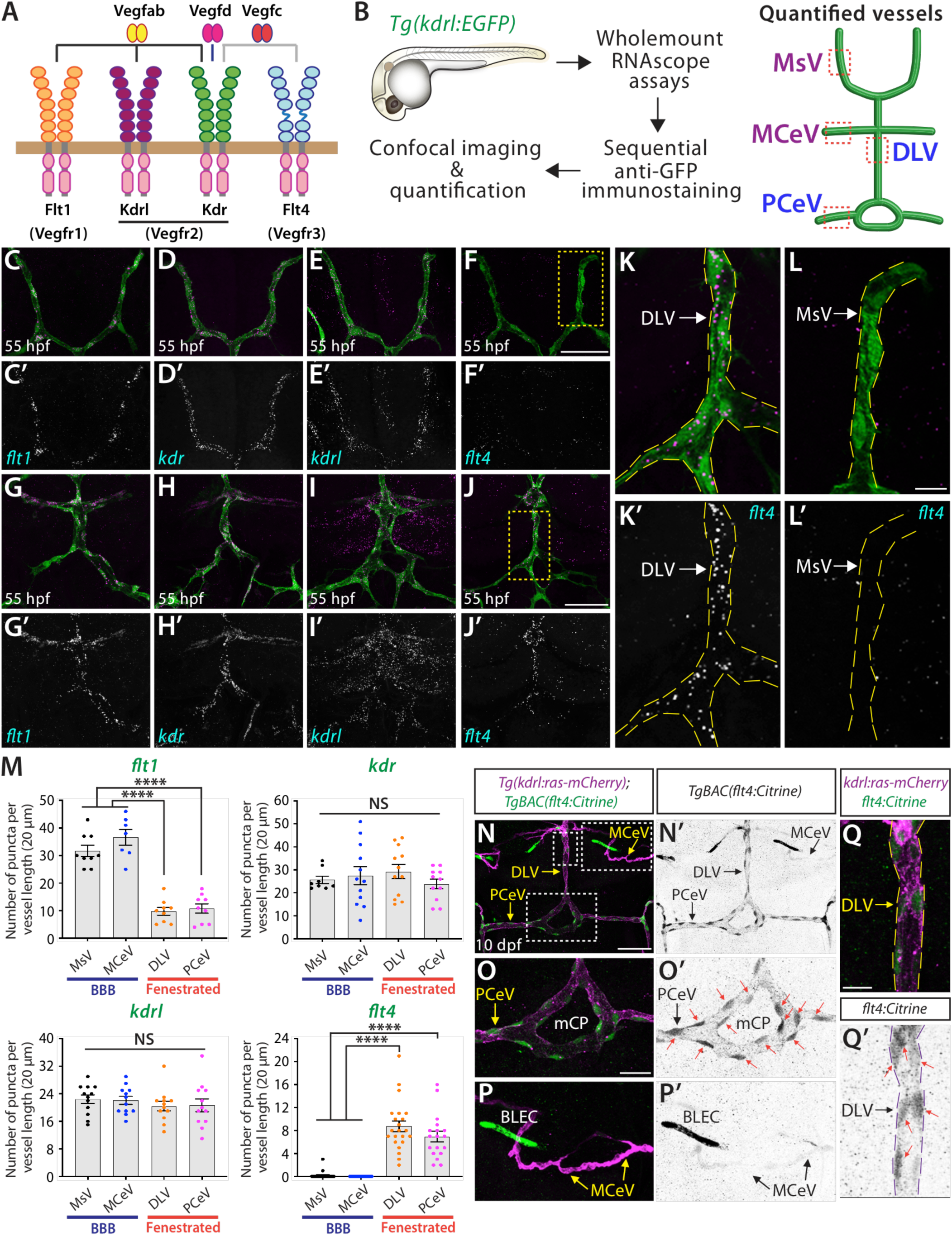
RNAscope analysis of *vegfr* expression during mCP vascularization. (**A**) Predicted receptor binding partners of Vegfab, Vegfc, and Vegfd in zebrafish. (**B**) Experimental setup for combined wholemount RNAscope, immunostaining, imaging, and quantification. Diagram shows the approximate vessel locations used for quantification. (**C**-**L’**) RNAscope assays in 55 hpf *Tg(kdrl:EGFP)* embryos using *flt1* (**C** and **G**), *kdr* (**D** and **H**), *kdrl* (**E** and **I**), and *flt4* (**F** and **J**) probes. Panels (**C**-**F’**) show the MsV, and panels (**G**-**J’**) show the DLV, PCeV, and MCeV. (**K**, **K’**) Magnified images from (**J**), showing *flt4* expression in the DLV and PCeV. (**L**, **L’**) Magnified images from (**F**), showing absent *flt4* expression in the MsV. (**M**) Quantification of mRNA signal per vessel in 55 hpf *Tg(kdrl:EGFP)* embryos for the indicated probes. (**N**, **N’**) Dorsal views of 10 dpf *TgBAC(flt4:Citrine);Tg(kdrl:ras-mCherry)* larval head, showing *flt4* reporter expression in the DLV and PCeV forming mCP vasculature. (**O**-**Q’**) Magnified views of the mCP region (**O**, **O’**), the MCeV (**P**, **P’**), and the DLV (**Q**, **Q’**) from (**N**). Robust *flt4* expression was detected in mCP vessels (red arrows), but not in the MCeV (arrows, **P’**). Scale bars: 50 µm in **F** for **C**-**F’**; 50 µm in **J** for **G**-**J’**; 10 µm in **L** for **K**-**L’**; 50 µm in **N** for **N’**; 20 µm in **O** for **O**-**P’**; 10 µm in **Q** for **Q’**.

To assess *vegfr* expression during cerebrovascular development, we performed RNAscope ISH at 55 and 75 hpf in EC-specific *Tg(kdrl:EGFP)* embryos (Fig. 3B). To compare expression levels between vessels, we quantified mRNA puncta per vessel length. At 55 hpf, both *kdr* and *kdrl* were uniformly expressed across dorsal cranial vessels (Fig. 3D-E’, H-I’, M). In contrast, we found that *flt4* expression was restricted to the DLV and PCeV (Fig. 3F-F’, J-L’, M), while *flt1* was more prominent in the MsV and MCeV (Fig. 3C-C’, G-G’, M). These overall expression patterns persisted at 75 hpf (Fig. S3A-K), identifying *flt4* expression as an early molecular distinction between sprouting mCP vessels and adjacent BBB vasculature, while also revealing all *vegfr* expression in developing mCP vessels.

### Brain vessel-selective *flt4* expression persists at later developmental stages

To further validate brain vessel-selective *flt4* expression, we utilized the established BAC reporter line *TgBAC(flt4:Citrine)* (*36*). This reporter strongly labeled lymphatic-like ECs in the primitive meninx during larval stages, which were previously identified as brain lymphatic EC (BLEC) (*37*). Although less pronounced than its robust expression in BLEC, *flt4* reporter activity was detected in the DLV, PCeV, and mCP vasculature at 10 dpf (Fig. 3N-Q’), consistent with RNAscope results at the same stage (Fig. S3L-P). These findings demonstrate *flt4*’s specific and persistent expression in mCP vessels and indicate that Flt4 signaling contribute to the development of heterogeneous vascular networks in this brain region through the regulation of mCP vasculature.

### Co-requirements of Vegfr2 and Vegfr3 signaling for mCP vascularization

To investigate the functional roles of each Vegfr in mCP vascularization, we analyzed their null mutants (Fig. 4A-F). Consistent with previous reports, the genetic loss of *kdrl* severely disrupted BBB angiogenesis, including failure of the central artery to invade the brain parenchyma (*9, 38*). Despite this severe defect, mCP vascularization remained intact, suggesting a distinct role for Kdrl signaling in BBB angiogenesis versus mCP vascularization. Similarly, other *vegfr* individual knockouts did not display mCP vascularization deficits.

**Fig. 4.**
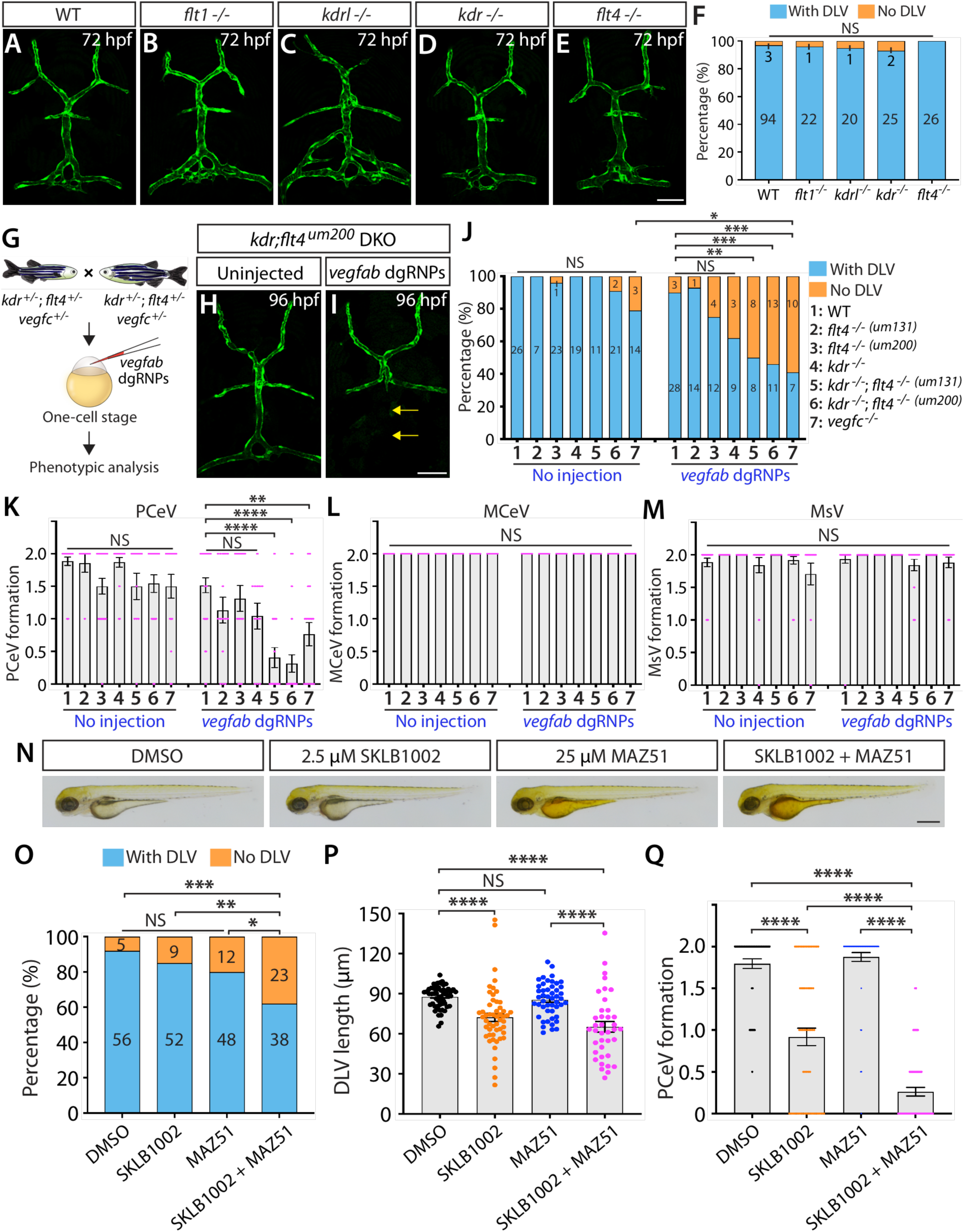
Genetic and pharmacological inhibition of Vegfr2 and Vegfr3 signaling exacerbated mCP vascularization defects. (**A**-**E**) Dorsal views of 72 hpf WT (**A**), *flt1^-/-^* (**B**), *kdrl^-/-^* (**C**), *kdr^-/-^* (**D**), and *flt4^-/-^* (**E**) cranial vasculature in *Tg(kdrl:EGFP)* larvae. (**F**) Percentage of larvae with and without the DLV for each genotype (number of animals examined per genotype is listed). (**G**) Experimental workflow for microinjection experiment (**H**-**M**). (**H** and **I**) 96 hpf *Tg(kdrl:EGFP)* larval images after no injection (**H**) and *vegfab* dgRNP injections (**I**). (**J**) Percentage of 96 hpf larvae of indicated genotypes and treatments with and without the DLV (number of animals examined per genotype and treatment is listed). (**K**-**M**) Quantification of PCeV (**K**), MCeV (**L**), and MsV (**M**) formation in uninjected and *vegfab* dgRNPs injected larvae at 96 hpf. (**N**) Brightfield images of 72 hpf larvae after treatment with DMSO, 2.5 μM SKLB1002, 25 μM MAZ51, or both SKLB1002 and MAZ51 from 34 to 72 hpf. (**O**) Percentage of the 72 hpf *Tg(kdrl:EGFP)* larvae with and without the DLV after the indicated treatments (number of animals examined per treatment is listed). (**P**) DLV length quantification in 72 hpf *Tg(kdrl:EGFP)* larvae after the indicated treatments. (**Q**) Quantification of PCeV formation in 72 hpf *Tg(kdrl:EGFP)* larvae after the indicated treatments. Scale bars: 50 µm in **E** for **A**-**E**; 50 µm in **I** for **H**; 500 µm in **N**.

Next, we investigated the combined roles of these receptors, given their significant genetic interactions during vascular developmental (*33, 34, 39*). To efficiently screen potential Vegfr combinations, we used the streamlined F0 knockout approach described earlier (*27*). By analyzing *kdr*, *kdrl*, and *flt4* knockouts in all possible combinations, we found that the combined loss of *flt4* and either *kdr* or *kdrl* did not result in mCP vascularization defects, while deletions of both *kdr* and *kdrl* caused severe embryonic vascular defects consistent with prior studies (*34, 39*). Despite these global angiogenic disruptions, we frequently observed DLV-like vessel formation, suggesting that these Vegfr2 paralogs are not the only receptors required for DLV formation.

To explore a potential role for Flt4 in this process, we induced the genetic loss of *kdr* and *flt4* in *vegfab* knockouts. This strategy was expected to mitigate global angiogenic defects caused by *kdrl* deletion, while reducing this receptor signaling dependent on Vegfab activity. To validate this approach, we injected a cocktail of *vegfab* dgRNPs into F0 progeny from *vegfc^+/-^* incrosses and analyzed phenotypes at 96 hpf (Fig. 4G). In *vegfc^-/-^*larvae injected with *vegfab* dgRNPs, we observed a significant increase in DLV loss phenotypes compared to uninjected *vegfc^-/-^* and injected *vegfc^+/+^* animals (Fig. 4J), recapitulating the co-requirement of Vegfab and Vegfc for DLV formation (*18*).

Using this approach, we next assessed mCP vascularization in single and double mutants of *kdr* and *flt4* with and without *vegfab* dgRNP injections. While *vegfab* dgRNPs-injected *kdr* and *flt4* single mutants did not exhibit significantly increased defects in DLV and PCeV formation, *kdr;flt4* double mutants displayed vascular defects comparable to those observed in injected *vegfc^-/-^* larvae (Fig. 4J-K). Notably, these injected *kdr;flt4* double mutants showed no defects in MsV and MCeV formation (Fig. 4L-M). These findings suggest a co-requirement of Kdr, Flt4, and Kdrl in mCP vascularization.

### Genetic mutants lacking the Flt4 cytoplasmic domain phenocopy *flt4* null mutant mCP vascularization defects

To dissect Flt4 function in mCP vascularization, we utilized the *flt4^um200^*allele, which lacks tyrosine residues in the cytoplasmic domain necessary for activating PI3K and MEK/ERK pathways (*40*). This mutant allowed us to assess the role of specific Flt4 cytoplasmic signaling in this process.

Using the same experimental workflow as in earlier analyses, we obtained results in *flt4^um200^* mutants that are consistent with those from the null *flt4^um131^* allele. Specifically, *flt4^um200^* single mutants did not show defects in DLV and PCeV formation, regardless of *vegfab* dgRNP injections. Conversely, *kdr;flt4^um200^*double mutants displayed severe vascularization defects (Fig. 4H-I), comparable to those observed in injected *vegfc^-/-^* larvae and *kdr;flt4^um131^* double mutants (Fig. 4J-M). These findings indicate that Flt4-dependent activation of the PI3K and MEK pathways is critical for mCP vascularization.

### Chemical co-inhibition of Vegfr2 and Flt4 signaling exacerbates mCP vascularization defects

To overcome the limitations of the *kdr;kdrl* double mutant analysis, which led to global angiogenic defects, we employed a pharmacological approach. Specifically, we used SKLB1002 and MAZ51 as Vegfr2- and Flt4-specific inhibitors, respectively, to treat embryos between 34 and 72 hpf. SKLB1002 treatment inhibited DLV extension in a dose-dependent manner (Fig. S4A-I) and reduced PCeV growth without causing overt morphological abnormalities (Fig. 4N, 4Q). However, SKLB1002 alone did not significantly increase DLV-absent phenotypes, regardless of dosage (Fig. 4O). Conversely, MAZ51 treatment alone caused a slight increase in DLV-absent phenotypes (Fig. 4O), but did not significantly affect DLV extension or PCeV formation (Fig. 4P-Q). Strikingly, co-treatment with both inhibitors dramatically exacerbated DLV-absent phenotypes and PCeV formation defects (Fig. 4N-Q), demonstrating a synergistic requirement of Vegfr2 and Flt4 signaling in these processes. The additional loss of *flt1* had no effect on these defects (Fig. S4J-M), ruling out Flt1 receptor signaling in this process. Collectively, both genetic and pharmacological results suggest distinct and redundant roles for Vegfr2 and Flt4 in mCP vascularization. Specifically, Flt4 and Vegfr2 paralogs are necessary for initial DLV sprouting, with Flt4 playing a more predominant role in this step. In contrast, Vegfr2 paralogs are primarily involved in extending the DLV and PCeV after sprouting by acting synergistically with Flt4.

## DISCUSSION

In this study, we address a long-standing question regarding the origins of cerebrovascular diversity. Through single-cell-resolution EC tracking at the border between fenestrated mCP and adjacent BBB vasculature, we uncover a previously uncharacterized niche within the MCeV that generates both EC subtypes. This discovery identifies the MCeV as a heterogeneous structure analogous to the lymphatic niche in the cardinal vein (*41*) and the haematopoietic niche in the aortic endothelium (*42–44*).

Our mechanistic characterization of this niche begins to elucidate how ECs are pre-programmed for distinct fates and initiate segregated capillary networks. Heterogeneous *glut1* expression in EC pools near the DMJ indicates pre-determination of BBB and fenestrated fates prior to angiogenesis. This Glut1 expression in pre-migratory ECs is reminiscent of BBB marker induction in ECs primed to invade the brain parenchyma (*10, 24, 45*). Additionally, we identified restricted *flt4* expression that distinguishes sprouting DLV and PCeV from adjacent BBB vessels, with this expression lasting until at least 10 dpf. This prolonged expression contrasts with transient *flt4* expression observed in tip cells during sprouting angiogenesis (*46, 47*) and aligns with earlier findings of VEGFR3 expression in fenestrated capillaries of the fetal and adult human CP (*48*). In mature vasculature, Flt4 is typically recognized as a lymphatic EC-specific marker, along with Prox1, Lyve1, and related genes (*49–52*). Given the emerging concept of hybrid vasculature expressing both blood and lymphatic identity markers (*53–57*), it will be interesting to investigate lymphatic gene expression in mCP vessels and uncover the upstream mechanisms driving selective *flt4* expression in brain capillaries.

To date, very few specific markers for fenestrated brain ECs have been identified. PLVAP has served as a primary marker distinguishing fenestrated ECs from BBB-associated ones in mature vasculature (*17, 58, 59*). Given the broad expression of PLVAP in immature ECs (*24, 59*), *flt4* emerges as a reliable early marker for fenestrated brain ECs. Moreover, we provide genetic and pharmacological evidence supporting Flt4’s vital function in regulating mCP vessel formation. Our observations of comparable angiogenic defects observed in two *flt4* mutant alleles demonstrate a requirement of this receptor signaling in this process. Notably, *flt4* mutants alone did not exhibit mCP vascularization defects, indicating that this process is independent of lymphatic formation. This finding is consistent with our prior observation of severe lymphatic defects in *vegfc;vegfd* double mutants with no disruption in mCP vascularization (*18*). Inhibitor treatments complement our genetic findings, showing that Vegfr2 and Flt4 cooperate in mCP vascularization by activating PI3K and ERK pathways as their major downstream effectors.

In summary, our study provides the first detailed characterization of a BBB and fenestrated vascular niche in the vertebrate brain, proposing a model in which Flt4-dependent signaling drives selective sprouting of fenestrated vessels. Our findings present a novel concept in brain vascularization, where distinct EC identities arise from specialized angioblasts in a local vascular niche through sequential processes of fate pre-determination and angiogenic segregation.

## Abbreviations

BBB: Blood-brain barrier
CP: Choroid plexus
CVOs: Circumventricular organs
dCP: Diencephalic choroid plexus
DLV: Dorsal longitudinal vein
DMJ: Dorsal midline junction
EC: Endothelial cell
Glut1: Glucose transporter 1
MCeV: Middle cerebral vein
MCtA: Mesencephalic central artery
mCP: Myelencephalic choroid plexus
MsV: Mesencephalic vein
MtA: Metencephalic artery
PCeV: Posterior cerebral vein
Plvap: Plasmalemma vesicle-associated protein
Vegfs: Vascular endothelial growth factors
Vegfrs: Vascular endothelial growth factor receptors

## Acknowledgements

We thank Drs. Nathan Lawson, Saulius Sumanas, and Michael Taylor for kindly providing us with fish lines; Don Zeisloft and his team for zebrafish care and husbandry; Dr. Judith Drazba and her team for confocal and electron microscopy imaging; and all members of the Matsuoka laboratory for discussions and comments on the manuscript. This work was supported by funding from the National Institutes of Health (R01 NS117510) and start-up funds from the Cleveland Clinic Foundation to R.L.M.

## Data and materials availability

All data supporting the findings of this study are included within this manuscript. Correspondence and requests for materials should be addressed to Ryota L. Matsuoka.

## Materials and Methods

### Zebrafish husbandry and strains

All zebrafish husbandry was performed under standard conditions in accordance with institutional and national ethical and animal welfare guidelines. All zebrafish work was approved by the Cleveland Clinic’s Institutional Animal Care and Use Committee under the protocol number 00003285. The following lines were used in this study: *Tg(kdrl:EGFP)^s843^* (*60*); *Tg(kdrl:Has.HRAS-mcherry)^s896^* (*61*), abbreviated *Tg(kdrl:ras-mCherry)*; *TgBAC(flt4:Citrine)^hu7135^* (*36*); *KI(etv2-2A-Gal4)^ci32Gt^* (*26*); *Tg(UAS:Kaede)^rk8^* (*62*); *Tg(glut1b:mCherry)^sj1^* (*24*), abbreviated *Tg(glut1:mCherry)*; *Tg(plvapb:EGFP)^sj3^* (*24*), abbreviated *Tg(plvap:EGFP)*; *vegfc^hu6410^* (*63*); *flt1^bns29^* (*35*); *kdrl ^um19^* (*64*); *kdr^bns32^* (*65*); *flt4^um131^*(*66*); and *flt4^um200^* (*40*). Adult fish were maintained on a standard 14 h light/10 h dark daily cycle. Fish embryos/larvae were raised at 28.5°C. To prevent skin pigmentation, 0.003% phenylthiourea (PTU) was used beginning at 10-12 hpf for imaging. Fish larvae analyzed at 10 dpf were transferred to a tank containing approximately 250 mL water supplemented with 0.003% PTU (up to 25 larvae/tank) and fed with Larval AP100 (<50 microns dry diet, Zeigler) starting at 5 dpf.

### Genotyping of mutants

Genotyping of *flt1^bns29^* mutant fish was performed by high-resolution melt analysis (HRMA) of PCR products as described previously (*35*). Genotyping of *kdrl ^um19^*, *kdr^bns32^*, and *flt4^um131^* mutant fish was performed by HRMA of PCR products using the following primers:

*kdrl um19* forward: 5’ – TGCTTCCTGATGGAGATACACACC – 3’

*kdrl um19* reverse 5’ – TGCAAATGAGTGTGAGTGTCCCAC – 3’

*kdr bns32* forward: 5’ – GACCTCACCCTGAGTCCACA – 3’

*kdr bns32* reverse: 5’ – GCGGTGCAGTTGAGTATGAG – 3’

*flt4 um131* forward: 5’ – GACCATCTTCATAACAGACTCTG – 3’

*flt4 um131* reverse: 5’ – GGATCTGAAACCAGACATGGTAC – 3’

Genotyping of *flt4^um200^*mutant fish was performed by PCR using the following primers:

*flt4 um200* forward: 5’ – GAAATCCCGTTCAATGTGTCTCAG – 3’

*flt4 um200* reverse: 5’ – CATTTCCATAGGAAGTTCCTCAAAG – 3’

*flt4 um200* forward: 5’ – acgtagccttatataactaagccc – 3’

The PCR reaction yielded products of 459 bp for the WT allele and 144 bp for the mutant allele.

### High-resolution melt analysis (HRMA)

A CFX96 Touch Real-Time PCR Detection System (Bio-Rad) was used for the PCR reactions and subsequent HRMA. Precision melt supermix for high-resolution melt analysis (Bio-Rad) was used in these experiments. PCR reaction protocols were: 95°C for 2 min, 46 cycles of 95°C for 10 s, and 60°C for 30 s. Following the PCR, a high-resolution melt curve was generated by collecting EvaGreen fluorescence data in the 65–95°C range. The analyses were performed on normalized derivative plots.

### Generation of F0 knockouts

Dual-guide ribonucleoprotein complexes (dgRNPs) consisting of CRISPR RNA (crRNA), trans-activating RNA (tracrRNA), and Cas9 protein were prepared as previously described (*27*). The following sequences of *vegfab* and *vegfc* crRNAs were used:

*vegfab* crRNA1: 5’ – AGTGCCTACATACCCAGAGA – 3’

*vegfab* crRNA2: 5’ – TTGTTATACACCTCCATGAA – 3’

*vegfab* crRNA3: 5’ – GAAGCGTCACAATAAATAAC – 3’

*vegfc* crRNA1: 5’ – CGTGACTTGACTCGAAAGCA – 3’

*vegfc* crRNA2: 5’ – CTGAACGCAACTGCTCCACC – 3’

*vegfc* crRNA3: 5’ – AAGTAAGCGCGACCCCATCT – 3’

Injection cocktails (4 μL) containing both *vegfab* and *vegfc* dgRNPs were prepared as follows: 0.4 μL 25 μM *vegfab* crRNA1:tracrRNA, 0.4 μL 25 μM *vegfab* crRNA2:tracrRNA, 0.4 μL 25 μM *vegfab* crRNA3:tracrRNA, 0.4 μL 25 μM *vegfc* crRNA1:tracrRNA, 0.4 μL 25 μM *vegfc* crRNA2:tracrRNA, 0.4 μL 25 μM *vegfc* crRNA3:tracrRNA, 1.2 μL 50 μM Cas9 protein stock, and 0.4 μL 0.5% phenol red solution (#P0290, Sigma). Injection cocktails (4 μL) containing only *vegfab* dgRNPs were prepared as follows: 0.4 μL 25 μM *vegfab* crRNA1:tracrRNA, 0.4 μL 25 μM *vegfab* crRNA2:tracrRNA, 0.4 μL 25 μM *vegfab* crRNA3:tracrRNA, 1.2 μL 25 μM Cas9 protein stock, 1.2 μL H_2_O, and 0.4 μL 0.5% phenol red solution. Injection cocktails were freshly prepared on the day of injection, and approximately 2 nL of the cocktails was injected into the cytoplasm of one-cell stage embryos.

### Immunohistochemistry

Immunohistochemistry was performed by following standard immunostaining procedures as described previously (*18*), except pErk immunostaining that is detailed below. fish were fixed in pH adjusted (pH 7.0) 4% paraformaldehyde (PFA)/phosphate buffered saline (PBS) overnight at 4°C and dehydrated through immersion in methanol serial dilutions (50%, 75%, then 100% methanol three times, 10 min each) at room temperature (RT). The dehydrated samples were stored in 100% methanol at −20°C until use. The samples were rehydrated through immersion in methanol serial dilutions (50%, then 25% methanol, 10 min each) at RT and washed briefly three times in 1% PBST (1% Triton X-100 in 0.1M PBS) followed by permeabilization in 10 µg/ml Proteinase K in 1% PBST at RT for 15 min. Samples were blocked at RT for 2-4 h in the blocking solution containing 0.5% bovine serum albumin in 1% PBST prior to primary antibody incubation.

Antigen retrieval of pErk immunostaining was performed as follows. After rehydration, samples were briefly equilibrated in the antigen retrieval buffer (150 mM Tris-HCl, pH 9.0) at RT for 5 min. The samples were then heated in this buffer at 70°C for 15 min. Samples were washed three times in 1% PBST at RT for 5 min each and permeabilized in 10 µg/ml Proteinase K in 1% PBST at RT for 15 min. The samples were then processed in the same manner as described above.

The following primary antibodies were used: chicken anti-GFP (GFP-1010, Aves Labs, 1:1000), rabbit anti-DsRed (#632496, Clontech, 1:300), and rabbit anti-Phospho-ERK1/2 (#4370S, Cell Signaling, 1:250). After primary antibody incubation at 4°C overnight, all samples were washed, incubated with secondary antibodies, and processed for imaging, as described previously (*18*).

### RNAscope *in situ* hybridization

RNAscope assays were performed on *Tg(kdrl:EGFP)^s843^*embryos and larvae collected at 55 hpf, 75 hpf, and 10 dpf. Samples were fixed, dehydrated, and rehydrated as detailed in the immunohistochemistry procedure. Permeabilization was performed using Protease Plus (#322331; Advanced Cell Diagnostics, ACD) at 40°C in the HybEZ^TM^ II hybridization system (#321711, ACD) for 5 minutes, 10 minutes, and 15 minutes for 55 hpf, 75 hpf, and 10 dpf samples, respectively. After permeabilization, samples were washed three times in 1% PBST at RT and transferred to 2 mL Eppendorf tubes (7-8 fish per tube). Negative control and gene-specific probes were designed and synthesized by ACD and prepared according to their instructions. To optimize this wholemount protocol for zebrafish samples, a probe targeting the bacterial gene *dapB* (#320871, 1X) was used as a negative control and confirmed to produce no detectable background signals. Probes specific to *flt1* (#815251-C3, 50X), *kdrl* (#416611-C2, 50X), and *flt4* (#1205121-C2, 50X) were freshly diluted in probe diluent (#300041, ACD) for each assay. A *kdr*-specific probe (#1318021-C1, 1X) was used without dilution. Hybridization with each probe was performed at 40°C for 2 hours. During this incubation, 1X wash buffer was diluted from the 50X stock (#310091, ACD) in H_2_O and pre-warmed.

Following hybridization, signal detection was carried out using the RNAscope Multiplex Fluorescent Detection Kit v2 (#323110, ACD). Samples were washed three times in pre-warmed wash buffer at 40°C for 10 min each. After this step, pre-warmed amplification reagents (Amp1, Amp2, Amp3) were sequentially applied at 40°C for 30 min, 30 min, and 15 min, respectively, with two intervening washes in wash buffer at RT for 5 min each. After incubation with Amp3, samples were washed two times in wash buffer at RT for 5 min each, and kept in 5X SSC (750 mM NaCl, 75 mM sodium citrate, pH 7.0) at RT overnight.

The following day, after washing once in wash buffer for 5 min at RT, samples were incubated with pre-warmed appropriate HRP-conjugated reagents (HRP-C3, HRP-C2, or HRP-C1) at 40°C for 15 min followed by two washes in wash buffer at RT for 5 min. Fluorescent signal was developed using TSA Vivid Fluorophore 650 (#323273, ACD) diluted at 1:750 in Multiplex TSA buffer (#322810, ACD) at 40°C for 30 min. After this step, samples were protected from prolonged light exposure to preserve fluorescence. After washing twice in wash buffer at RT for 5 min each, samples were treated with pre-warmed Multiplex FL v2 HRP Blocker (#323107, ACD) at 40°C for 15 min, washed once in 1% PBST at RT, and incubated with anti-GFP antibody described earlier at RT for 6 hours. After three washes in 1% PBST at RT for 20 min each, samples were incubated with a secondary antibody at 4 °C overnight. After six washes in 1% PBST at RT for 15 min each, samples were processed for imaging.

### Quantification of RNAscope Assays

Gene expression in the DLV, PCeV, MsV, and MCeV was quantified using high-resolution images acquired from RNAscope assays. For 55 and 75 hpf samples, a 20 μm x 20 μm area was selected, and the number of puncta signals overlapping with *Tg(kdrl:EGFP)*-positive ECs was manually counted. The specific locations used for this quantification are indicated in the vascular diagram shown in Fig. 3B. For 10 dpf larvae, a 50 μm x 50 μm area was quantified at locations similar to those used for 55 and 75 hpf embryos/larvae. For probes with some background signals, only the bright signals were counted. For most probes, maximum projection z-stack images were used for signal counting, as these probes displayed highly specific vascular signals. For some probes with strong non-blood vessel signals, z-stack images were carefully checked stack-by-stack to count only signals overlapping with *Tg(kdrl:EGFP)*-positive ECs. The quantified data for all probes using this method well reflect our visual observations of the gene expression patterns.

### Photoconversion

Photoconversion experiments were performed in embryos carrying the *KI(etv2-2A-Gal4)^ci32Gt^* and *Tg(UAS:Kaede)^rk8^*transgenes using a Leica SP8 confocal microscope as follows. A dorsal view image was first captured for each embryo at approximately 43 hpf to select target cells for photoconversion. Using the bleach point setting in Leica Application Suite X (LAS X) software, a region of interest (ROI) was designated for each target cell. Either single or multiple cells were targeted per embryo. Photoconversion of Kaede from green to red fluorescence was achieved by applying 405 nm laser illumination (1% power) for 1 sec. If photoconversion appeared incomplete, an additional illumination for 1 sec was applied. After successful photoconversion, bleach point setting was deactivated, and post-photoconversion images were captured. Photoconversion in *vegfab;vegfc* crispant embryos was performed in the same manner, except that these embryos were injected at the one-cell stage with a cocktail containing both *vegfab* and *vegfc* dgRNPs.

For longitudinal cell tracking, photoconverted embryos were carefully removed from agarose gels using a glass pipette, transferred to 24-well plates containing PTU egg water, and maintained at 28.5°C. Images were captured approximately 24 hours later at 67 hpf to assess migration destinations of photoconverted cells. For time-lapse imaging, embryos were imaged immediately after photoconversion in a temperature-controlled chamber maintained at 28.5°C. Images were acquired at 8-10 min intervals from approximately 43 to 53 hpf to monitor cell migration dynamics.

### Quantification of photoconverted cell destinations

To quantify the migration destinations of photoconverted cells, their initial positions at 43 hpf were measured relative to the midline using a defined ROI. These positions were categorized based on their distances from the midline on both hemispheres, as shown in Fig. 1N. Cells located within 50 µm from the midline were assigned to the 1st position, while those located between 50 and 100 µm were assigned to the 2nd position. When multiple targeted cells within an embryo were distributed across both positions, data from these embryos were excluded from the graphs shown in Fig. 1N, but were included in the graph presented in Fig. 1T. Migration destinations were determined by the positions of photoconverted cell bodies at 67 hpf and categorized into five locations: DLV/PCeV (mCP vessels), MCtA, MsV, MsV-DLV junctions, and MCeV

### Confocal and stereo microscopy

Fluorescence imaging of live and fixed zebrafish samples was conducted using a Leica TCS SP8 confocal laser scanning microscope. Fish embryos and larvae were anaesthetized with a low dose of tricaine, embedded in 1% low melt agarose in glass-bottom Petri dishes (MatTek), and imaged using a 25X water immersion objective lens. Image acquisition and analysis were conducted using LAS X software (Version 3.7.0.20979). Brightfield images of anesthetized fish were captured using a Nikon SMZ-18 stereomicroscope. Image acquisition and analysis for brightfield imaging were carried out using NIS-Elements BR Imaging software (Version 5.10.01).

### Pharmacological treatment

Embryos carrying the *Tg(kdrl:EGFP)^s843^* transgene were used for pharmacological experiments. Treatments were conducted from 34 to 72 hpf with the following inhibitors: SL327 (#S4069, Sigma), LY294002 (#L9908, Sigma), U73122 (#U6756, Sigma), SKLB1002 (#SML1164, Sigma), and MAZ51 (#676492, Sigma). At 54 hpf, each drug solution was replaced with a fresh preparation to ensure consistent efficacy of the chemicals. All chemicals were initially prepared as 100X stock solutions dissolved in 100% DMSO and diluted 1:100 in egg water immediately before use.

### Quantification of DLV, PCeV, MsV, and MCeV formation

Quantification was performed using fish carrying the *Tg(kdrl:EGFP)^s843^* reporter. Fish larvae at the indicated developmental stages were analyzed for the presence or absence of the DLV. In larvae where the DLV was present, the lengths of the DLV were measured by drawing a ROI using the polyline tool in LAS X software. To assess the bilateral formation of the PCeV, MsV, and MCeV, the following scoring criteria were applied: 1) Score 2 - both bilateral vessels are fully formed; 2) Score 1.5 - one vessel is fully formed, while the other is partially formed; 3) Score 1 - both vessels are partially formed, or one vessel is fully formed while the other is absent; 4) Score 0.5 - one vessel is partially formed, while the other is absent; and 5) Score 0 - both bilateral vessels are absent.

### Statistical analysis

Statistical differences in mean values among multiple groups were determined using a one-way analysis of variance (ANOVA) followed by Tukey’s multiple comparison test. Fisher’s exact test was applied to assess significance when comparing the degree of penetrance of observed phenotypes. Statistical analyses were conducted using GraphPad Prism 8.1.1. The criterion for statistical significance was set at *P* < 0.05. Error bars represent SEM.

