## Supplemental Materials for "Angiogenic mechanisms governing the segregation of blood-brain barrier and fenestrated capillaries derived from a multipotent cerebrovascular niche"

### Figure S1

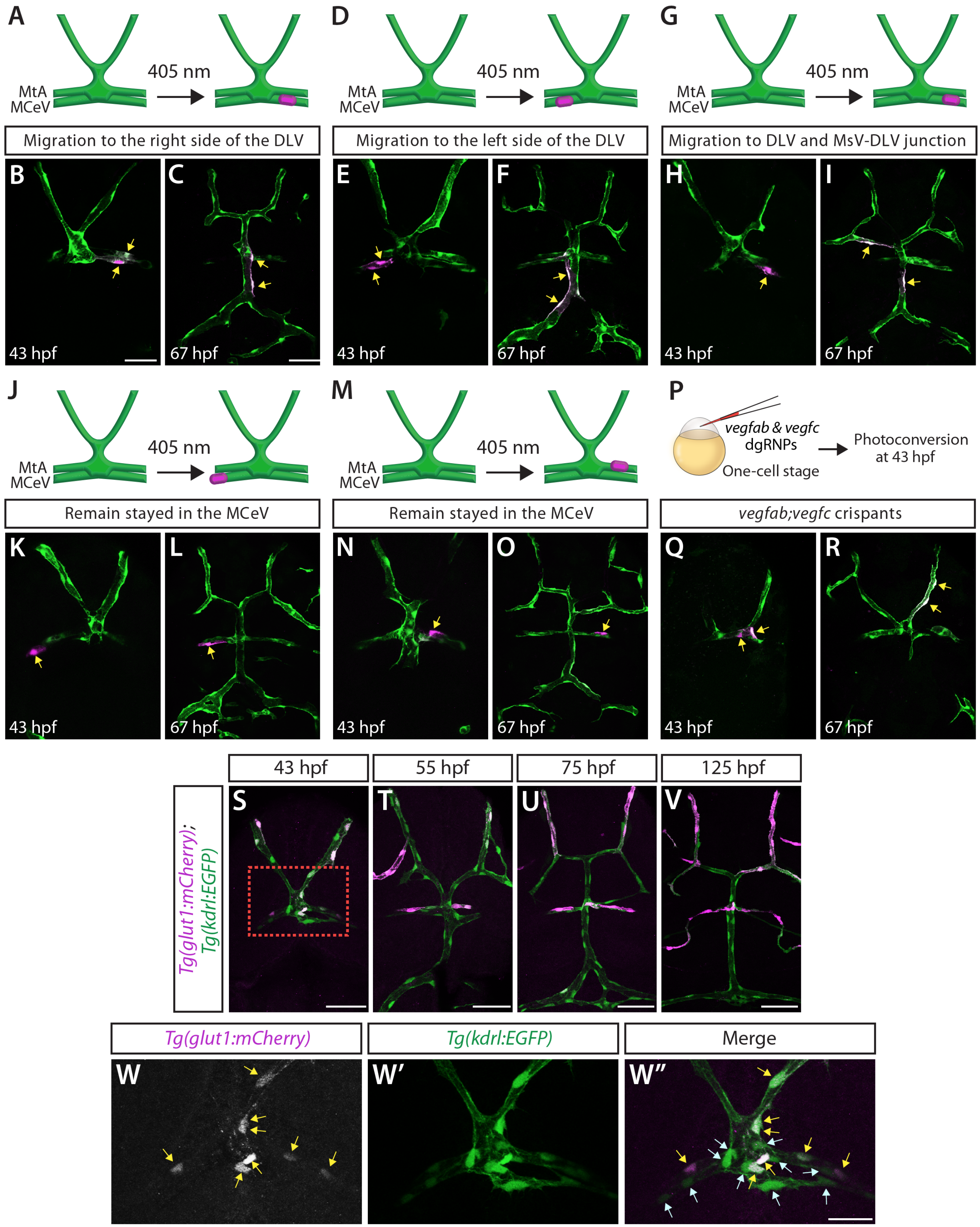

A
405 nm
MtA
MCeV
Migration to the right side of the DLV
D
405 nm
MtA
MCeV
Migration to the left side of the DLV
G
405 nm
MtA
MCeV
Migration to DLV and MsV-DLV junction
B
43 hpf
C
67 hpf
E
43 hpf
F
67 hpf
H
43 hpf
I
67 hpf
J
405 nm
MtA
MCeV
Remain stayed in the MCeV
M
405 nm
MtA
MCeV
Remain stayed in the MCeV
P
vegfab & vegfc dgRNPs
One-cell stage
Photoconversion at 43 hpf
vegfab;vegfc crispants
K
43 hpf
L
67 hpf
N
43 hpf
O
67 hpf
Q
43 hpf
R
67 hpf
Tg(glut1:mCherry); Tg(kdrl:EGFP)
43 hpf
55 hpf
75 hpf
125 hpf
S
T
U
V
Tg(glut1:mCherry)
Tg(kdrl:EGFP)
Merge
W
W'
W''

**Fig. S1. Photoconversion tracking from the MCeV and MtA, and heterogeneous *glut1* expression at the DMJ.**

(**A**, **D**, **G**, **J**, **M**) Diagrams indicating photoconversion locations for panels (**B**-**C**, **E**-**F**, **H**-**I**, **K**-**L**, and **N**-**O**, respectively).

(**B**, **C**) Photoconversion in 43 hpf *KI(etv2-2A-Gal4);Tg(UAS:Kaede)* embryo (**B**), with subsequent tracking at 67 hpf (**C**). Photoconverted cells in the right MCeV migrated to the right DLV (yellow arrows).

(**E**, **F**) Tracking of photoconverted cells from 43 hpf (**E**) to 67 hpf (**F**). Photoconverted cells in the left MCeV migrated to the left DLV (yellow arrows).

(**H**, **I**) Tracking of photoconverted cells from 43 hpf (**H**) to 67 hpf (**I**). Photoconverted cells in the right MCeV migrated to the two locations (yellow arrows): one on the right DLV and the other on the left MsV-DLV junction.

(**K**, **L**) Tracking of photoconverted cells from 43 hpf (**K**) to 67 hpf (**L**). A photoconverted cell in the left MCeV remained on the left MCeV (yellow arrows).

(**N**, **O**) Tracking of photoconverted cells from 43 hpf (**N**) to 67 hpf (**O**). A photoconverted cell in the right MtA remained on the same area (yellow arrows).

(**P**) Experimental setup for microinjection experiments (**Q**, **R**).

(**Q**, **R**) Tracking of photoconverted cells in *vegfab;vegfc* crispants from 43 hpf (**Q**) to 67 hpf (**R**). MCeV-derived photoconverted cells migrated to the MsV in crispants (yellow arrows).

(**S-W"**) Dorsal views of *Tg(kdrl:EGFP);Tg(glut1:mCherry)* heads immunostained for GFP and mCherry, showing expanded *glut1* reporter expression in the MsV and MCeV from 43 to125 hpf. Magnified views from (**S**) showed heterogeneous *glut1* expression in the DMJ (**W-W"**).

Scale bars: 50 µm in **B** for **E**, **H**, **K**, **N**, **Q**; 50 µm in **C** for **F**, **I**, **L**, **O**, **R**; 50 µm in **S-V**; 20 µm in **W-W"**.

### Figure S2

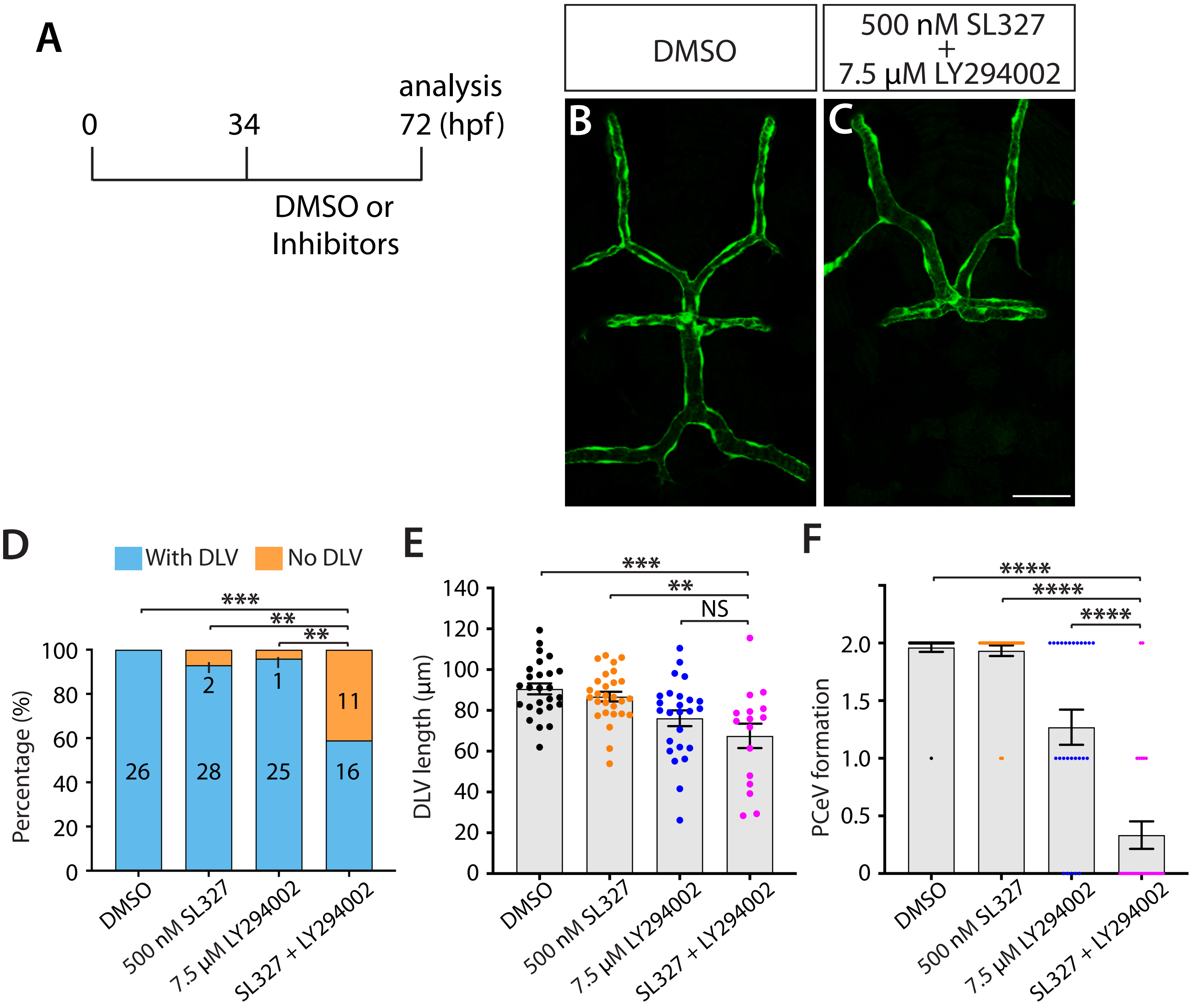


**Fig. S2. Co-inhibition of MEK and PI3K signaling exacerbated mCP vascularization defects.**
(**A**) Experimental timeline for chemical treatments (**B-H**).
(**B**, **C**) Dorsal views of 72 hpf *Tg(kdrl:EGFP)* larvae after treatment with DMSO (**B**) and a combined solution of 500 nM SL327 and 7.5 µM LY294002 (**C**), showing both DLV and PCeV formation defects with co-treatment.
(**D**) Percentage of 72 hpf *Tg(kdrl:EGFP)* fish with and without the DLV after the indicated treatment (number of animals examined per genotype is listed).
(**E**) DLV length quantification in 72 hpf *Tg(kdrl:EGFP)* larvae that formed the DLV (n=26 for DMSO, n=28 for 500 nM SL327, n=25 for 7.5 µM LY294002, and n=16 for combined SL327/LY294002).
(**F**) Quantification of PCeV formation in 72 hpf *Tg(kdrl:*EGFP) larvae after the indicated treatment (n=26 for DMSO, n=30 for 500 nM SL327, n=26 for 7.5 µM LY294002, and n=27 for combined SL327/LY294002).
Scale bars: 50 µm in **C** for **B**.

### Figure S3

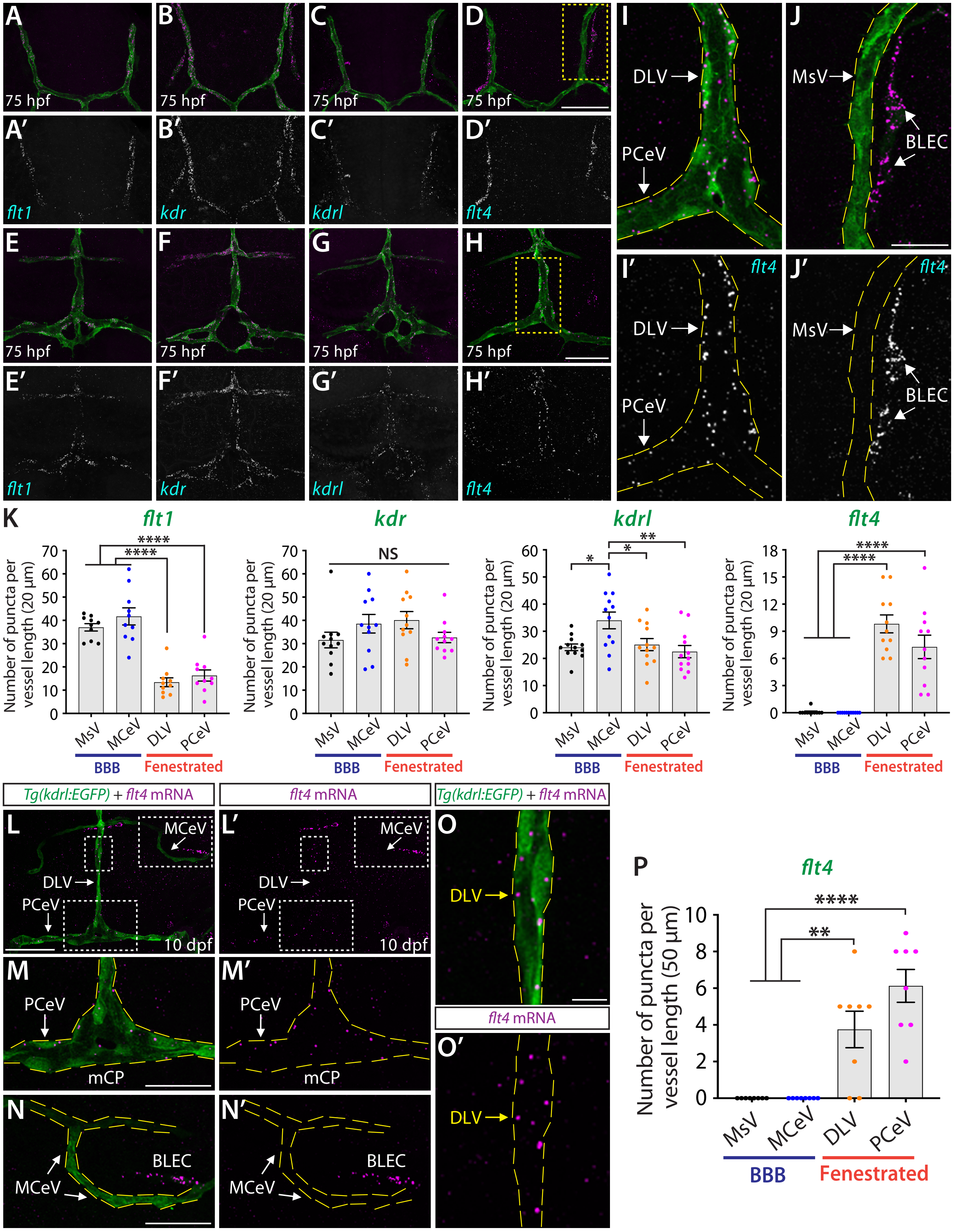

A
75 hpf
B
75 hpf
C
75 hpf
D
75 hpf
A′
flt1
B′
kdr
C′
kdrl
D′
flt4
E
75 hpf
F
75 hpf
G
75 hpf
H
75 hpf
E′
flt1
F′
kdr
G′
kdrl
H′
flt4
I
DLV
PCeV
J
MsV
BLEC
I′
flt4
DLV
PCeV
J′
flt4
MsV
BLEC
K
flt1
kdr
kdrl
flt4
Number of puncta per vessel length (20 μm)
NS
MsV
MCeV
DLV
PCeV
BBB
Fenestrated
Tg(kdrl:EGFP) + flt4 mRNA
flt4 mRNA
L
MCeV
DLV
PCeV
10 dpf
L′
O
M
PCeV
mCP
M′
N
MCeV
BLEC
N′
O′
P
flt4
Number of puncta per vessel length (50 μm)

**Fig. S3. Brain vessel-selective *flt4* expression during mCP vascularization and at 10 dpf**
(**A**-**J'**) RNAscope assays in 75 hpf *Tg(kdrl:EGFP)* larvae using *flt1* (**A** and **E**), *kdr* (**B** and **F**), *kdrl* (**C** and **G**), and *flt4* (**D** and **H**) probes. Panels (**A**-**D'**) show the MsV, and panels (**E**-**H'**) show the DLV, PCeV, and MCeV.
(**I**, **I'**) Magnified images from (**H**), showing *flt4* expression in the DLV and PCeV.
(**J**, **J'**) Magnified images from (**D**), showing absent *flt4* expression in the MsV.
(**K**) Quantification of mRNA signal per vessel in 75 hpf *Tg(kdrl:EGFP)* larvae for the indicated probes.
(**L**-**O'**) RNAscope assays in 10 dpf *Tg(kdrl:EGFP)* larvae using *flt4* probe. Magnified views of the mCP region (**M**, **M'**), the MCeV (**N**, **N'**), and the DLV (**O**, **O'**) from (**L**, **L'**).
(**P**) Quantification of *flt4* signal per vessel in 10 dpf *Tg(kdrl:EGFP)* larvae.
Scale bars: 50 µm in **D** for **A**-**D'**; 50 µm in **H** for **E**-**H'**; 20 µm in **J** for **I**-**J'**; 50 µm in **L** for **L'**; 25 µm in **M** for **M'**; 25 µm in **N** for **N'**; 10 µm in **O** for **O'**.

### Figure S4

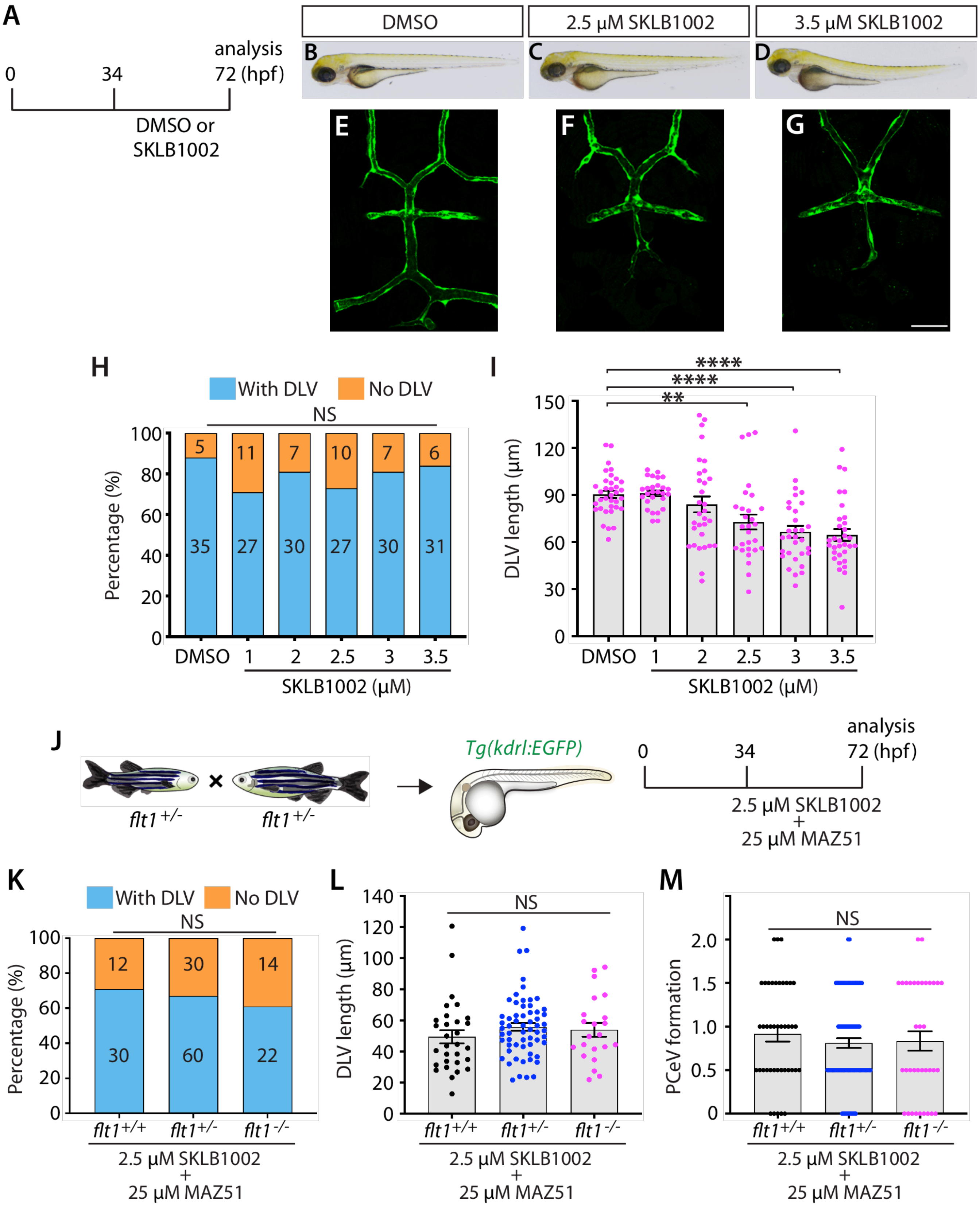

A
0
34
analysis
72 (hpf)
DMSO or
SKLB1002
DMSO
2.5 µM SKLB1002
3.5 µM SKLB1002
B
C
D
E
F
G
H
With DLV
No DLV
NS
Percentage (%)
DMSO
1
2
2.5
3
3.5
SKLB1002 (µM)
I
****
****
**
DLV length (µm)
DMSO
1
2
2.5
3
3.5
SKLB1002 (µM)
J
flt1+/-
×
flt1+/-
Tg(kdrl:EGFP)
0
34
analysis
72 (hpf)
2.5 µM SKLB1002
+
25 µM MAZ51
K
With DLV
No DLV
NS
Percentage (%)
flt1+/+
flt1+/-
flt1-/-
2.5 µM SKLB1002
+
25 µM MAZ51
L
NS
DLV length (µm)
flt1+/+
flt1+/-
flt1-/-
2.5 µM SKLB1002
+
25 µM MAZ51
M
NS
PCeV formation
flt1+/+
flt1+/-
flt1-/-
2.5 µM SKLB1002
+
25 µM MAZ51

**Fig. S4. Dose-dependent SKLB1002 effects on DLV formation, and comparison of mCP vessel formation in WT and *flt1* mutants treated with SKLB1002 and MAZ51.**

(**A**) Experimental timeline for chemical treatments (**B**-**G**).
(**B**-**D**) Brightfield views of 72 hpf *Tg(kdrl:EGFP)* larvae after treatment with DMSO (**B**), 2.5 μM SKLB1002 (**C**), and 3.5 μM SKLB1002 (**D**).
(**E**-**G**) Dorsal views of 72 hpf *Tg(kdrl:EGFP)* larval heads after treatment with DMSO (**E**), 2.5 μM SKLB1002 (**F**), and 3.5 μM SKLB1002 (**G**).
(**H**) Percentage of 72 hpf *Tg(kdrl:EGFP)* larvae with and without the DLV after the indicated SKLB1002 treatments (number of animals examined per genotype is listed).
(**I**) DLV length quantification in 72 hpf *Tg(kdrl:EGFP)* larvae that formed the DLV after SKLB1002 treatments (n=35 for DMSO, n=27 for 1 μM, n=30 for 2 μM, n=27 for 2.5 μM, n=30 for 3 μM, and n=31 for 3.5 μM).
(**J**) Experimental setup for chemical treatments (**K**-**M**).
(**K**) Percentage of 72 hpf larvae of the indicated genotypes with and without the DLV after co-treatment with 2.5 μM SKLB1002 and 25 μM MAZ51.
(**L**) DLV length quantification in 72 hpf fish of the indicated genotypes after SKLB1002 and MAZ51 co-treatment.
(**M**) Quantification of PCeV formation in 72 hpf larvae of the indicated genotypes after SKLB1002 and MAZ51 co-treatment.
Scale bars: 50 μm in **G** for **E**-**G**.

**Movie S1. Dynamic EC scrambling at the DMJ before and during DLV sprouting.**
Time lapse live imaging of a *Tg(kdrl:EGFP)* embryo from 32 hpf to 47 hpf, showing dynamic EC movements at the DMJ before and during DLV sprouting.

**Movie S2. Time-lapse imaging of a photoconverted MCeV cell migrating to the DLV.**
Tracking of a photoconverted MCeV cell in a *KI(etv2-2A-Gal4);Tg(UAS:Kaede)* embryo from 43 to 53 hpf, showing its migration trajectory to the DLV.
